# Histone H3 Ser10 phosphorylation occurs exclusively in replicative stages and peaks during mitosis in *Trypanosoma cruzi*

**DOI:** 10.64898/2026.02.16.706091

**Authors:** María del Rosario López, Salomé Catalina Vilchez Larrea, Josefina Ocampo, Guillermo Daniel Alonso

**Affiliations:** Instituto de Investigaciones en Ingeniería Genética y Biología Molecular “Dr. Héctor N. Torres” (INGEBI), Consejo Nacional de Investigaciones Científicas y Técnicas (CONICET), Buenos Aires, Argentina; Departamento de Fisiología, Biología Molecular y Celular, Facultad de Ciencias Exactas y Naturales, Universidad de Buenos Aires, Buenos Aires, Argentina

**Keywords:** H3Ser10 phosphorylation, Chromatin dynamics, Cell cycle regulation, Mitosis, Aurora kinase

## Abstract

Protein phosphorylation is a central post-translational modification that regulates signaling pathways across all living organisms. Through the antagonistic activities of protein kinases and phosphatases, phosphorylation modulates protein function by inducing conformational changes that affect enzyme activity, protein–protein interactions, stability, and subcellular localization. These molecular events regulate diverse cellular processes, including cell cycle progression, differentiation, gene expression, and metabolism. In unicellular parasites such as *Trypanosoma cruzi, Trypanosoma brucei*, and *Leishmania spp*., specialized signaling pathways have evolved to enable adaptation to the fluctuating environments of insect vectors and mammalian hosts. In many eukaryotes, phosphorylation of histone H3 at serine 10 (H3Ser10p) is essential for proper chromosome condensation during mitosis and is catalyzed by Aurora kinase B. Although trypanosomatids possess an Aurora kinase B homolog and a conserved serine residue at position 10 of histone H3, this modification had not been previously detected in these organisms. Here, using a stage-specific approach, we report the first detection of H3Ser10p in *T. cruzi* and explore its association with cell cycle progression. Western blot analyses using a specific antibody revealed H3Ser10p in exponentially growing epimastigotes, both in total protein extracts and nucleosome-enriched fractions, indicating its incorporation into chromatin. Fluorescence microscopy showed that this histone mark is restricted to the nuclei of dividing cells. Furthermore, H3Ser10p was detected exclusively in replicative stages of the parasite. Analysis of cell cycle–associated structures and flow cytometry demonstrated that H3Ser10 phosphorylation is dynamically regulated, peaking in the G2/M phase. These findings identify H3Ser10p as a novel epigenetic mark in *T. cruzi* that is tightly regulated during the cell cycle.

## Introduction

In eukaryotic cells, phosphorylation of serine 10 on histone H3 (H3Ser10p) is a highly conserved mitotic modification that appears at the onset of mitosis, specifically during prophase. This mark correlates with chromatin condensation, an essential process for accurate chromosome segregation and nuclear division (1,2). Beyond its structural role, H3Ser10p has been implicated in the maintenance of genome stability (3) and has been proposed to participate in checkpoint-like mechanisms that coordinate chromatin architecture with mitotic progression (4,5).

In multicellular eukaryotes, H3Ser10 phosphorylation is catalyzed by Aurora kinase B (AUKB), a core component of the chromosomal passenger complex (6). This complex orchestrates multiple mitotic events, including chromosome alignment, segregation, and cytokinesis, and its disruption leads to severe defects in cell division (7). Consistent with its pivotal role in cell cycle control, deregulation of H3Ser10p has been associated with genomic instability and oncogenic transformation (8,9). Owing to its strict mitotic restriction, H3Ser10p is widely used as a reliable biomarker for mitotic cells (10,11).

*Trypanosoma cruzi* exhibits a complex life cycle alternating between the insect vector and the mammalian host, involving profound morphological, metabolic, and proliferative changes. Notably, differentiation between life-cycle stages is tightly coupled to cell cycle regulation, as replicative and non-replicative forms coexist within the parasite’s developmental program. In trypanosomatids, while the overall structure of the cell cycle resembles that of other eukaryotes, several distinctive features are observed. These organisms undergo a closed mitosis in which the nuclear envelope remains intact (12), and although chromatin condensation occurs, classical mitotic chromosomes are not formed (13). Moreover, progression through the cell cycle requires the coordinated duplication and segregation of additional organelles, including the kinetoplast and the flagellum, adding further complexity to this process (14–16).

In trypanosomatids, regulation of the cell cycle is intimately linked to developmental transitions. In *Trypanosoma brucei*, cell cycle checkpoints have been shown to differ markedly from those of opisthokont eukaryotes, with differentiation often associated with cell cycle arrest or remodeling rather than canonical checkpoint activation (17). Epigenetic mechanisms play a central role in this process, as illustrated by the tight coupling between histone modifications and cell cycle progression. Notably, trimethylation of histone H3 at lysine 76 (H3K76me3), mediated by Dot1B, has been demonstrated to be essential for proper cell cycle control and differentiation in *T. brucei*, with its deregulation leading to aberrant DNA replication and developmental defects (18). These findings highlight the importance of dynamic histone modifications in coordinating proliferation and developmental competence in trypanosomes.

Trypanosomatids encode a small family of Aurora kinases (AUK1–3), among which AUK1 is the closest functional homolog of metazoan AUKB. In *T. brucei*, RNAi-mediated depletion of TbAUK1 results in severe cytokinesis defects, generation of polyploid cells with multiple kinetoplasts and flagella, and metaphase arrest, supporting a key role for this kinase in mitotic progression (19,20). In *T. cruzi*, TcAUK1 displays a dynamic subcellular localization during the cell cycle, localizing in kinetoplast-associated regions during G1/S and accumulating in the nucleus during G2/M. Perturbation of TcAUK1 expression affects cell cycle progression, leading to delays in G2/M, consistent with observations in *T. brucei* (21–23).

Despite the conservation of the N-terminal tail of histone H3 in trypanosomatids, including the serine residue at position 10, phosphorylation of H3Ser10 has not been previously reported in *T. cruzi* (24). Based on the evolutionary conservation of this residue, the nuclear localization of TcAUK1 during mitosis, and the central role of histone modifications in trypanosome cell cycle regulation and differentiation, we hypothesized that H3Ser10 phosphorylation could occur during mitosis in *T. cruzi* and contribute to chromatin regulation during cell division.

Here, we report for the first time the occurrence of H3Ser10 phosphorylation during mitosis in *Trypanosoma cruzi*. We show that H3Ser10p is associated with insoluble chromatin fractions, is detected in nucleosome core particle preparations, and is specifically removed by Lambda protein phosphatase treatment. Furthermore, H3Ser10p displays a strictly nuclear localization in mitotic cells and is observed exclusively in replicative life-cycle stages. Finally, we demonstrate that H3Ser10 phosphorylation is a dynamic, cell-cycle-regulated process, with increased signal intensity during mitosis, supporting its role as a bona fide mitotic histone modification in this parasite.

## Materials and methods

### *T. cruzi* epimastigotes culture

Epimastigotes of Tulahuen and Dm28c strains were grown at 28°C in LIT medium [5 g.l^-1^ liver infusion, 5 g.l^-1^ Bacto-tryptose, 68 mM NaCl, 5.3 mM KCl, 22 mM Na2PO4, 0.2% (w/v) glucose, 0.002% (w/v) hemin, containing 10% v/v fetal bovine serum (NATOCOR, Argentina), 100 units.ml^-1^ penicillin and 100 μg.l^-1^ streptomycin]. Cell density was maintained between 1×10^6^ and 1×10^8^ cells.ml^-1^. The Neubauer chamber was used for cell counting.

### Infection of Vero cells and obtention of trypomastigotes

*Cercopithecus aethiops* (green monkey) Vero cells (ATCC CCL-81) were cultured at 37°C and 5% CO2 supplied in Minimum Eagle Medium (MEM, Gibco) supplemented with 5% fetal bovine serum (NATOCOR, Argentina), 2 mM L-Glutamine (Sigma), 100 units.ml^-1^ penicillin and 100 μg.l^-1^ streptomycin.

For the obtention of *T. cruzi* trypomastigotes and amastigotes, Vero cells were infected with trypomastigotes (1:10 ratio) for 24 h, after which they were washed with Phosphate Buffered Saline (PBS: NaCl 137 mM, KCl 2.7 mM, Na2HPO4 10 mM, and KH2PO4 1.8 mM) and maintained in MEM-SFB 5%. Trypomastigotes in culture supernatant were harvested by centrifugation and processed as needed. Amastigotes were collected after 48 hours post infection (hpi) for different processes.

### Amastigotes intracellular purification

The protocol was done as previously reported (25). Briefly, infected Vero cells were detached by gently scraping them in 5 mL PBS and then passed thrice through a 27 ½ gauge needle connected to a 50 mL syringe. The liquid obtained from this process was centrifuged at 400 xg for 10 min to eliminate any cellular debris from the host cell. The supernatant was transferred to a sterile tube and centrifuged at 1000 xg for 10 min. Amastigotes were recovered within the pellet.

### Immunofluorescence and Immunolocalization

Epimastigotes were harvested by centrifugation at 1000 xg for 5 min, washed once with PBS, and fixed with 4% paraformaldehyde (PFA) for 20 min. Next, the samples were adhered to poly-L-lysine-coated coverslips. Vero cells were cultured in 24 wells plates with a sterile coverslip and infected with Dm28c strain. Forty-eight hpi, the culture medium was removed. Then, the cells were washed once with PBS before further processing and fixed as indicated for epimastigotes. After washing samples twice with PBS, they were permeabilized with PBT solution (0.2% Triton-×100 in PBS) and incubated in blocking solution (3% BSA in PBS) for 1 h at room temperature. To check the H3Ser10 phosphorylation state, a specific anti-H3Ser10p for this modification was used (ab177218, ABCAM) at 1:300, incubating for 1 h at room temperature. In parallel, anti-tubulin antibody (TS168, SIGMA) was used at 1:1000. The secondary antibodies (anti-Mouse AF546 and anti-Rabbit AF488, INVITROGEN) were incubated for 1 h at room temperature. Samples were mounted with VectaShield with DAPI (Vector Labs) and then visualized using an Olympus BX41/FV300. Images were processed with ImageJ software.

### MNase digestion of chromatin

*T. cruzi* epimastigotes of CL Brener strain were grown to log-phase in LIT medium. The cells were centrifuged for 5 min at 1000 xg and resuspended in fresh LIT medium before fixation. Cells were fixed with 1% formaldehyde (20 min at room temperature), and glycine was added to 0.4 M for 20 min with gentle agitation. Cells were centrifuged for 25 min at 3,200 xg at 4 °C. The pellet was stored at −80 °C until required. For MNase digestion, the pellet was resuspended in 700 µl of Complete Lysis Buffer (1 mM L-glutamine, 250 mM sucrose, 2.5 mM CaCl2, 1 mM phenylmethylsulphonyl fluoride (PMSF), 0.01% (v/v) Triton X 100) and divided into seven aliquots of 400 µL, which were digested with increasing amounts of MNase (Worthington). The digestion was stopped by adding 100 µL of Stop buffer (5 mM Na-EDTA, pH 7.5, 5 mM Na-EGTA, 0.05% (v/v) NP40). The extent of digestion was verified by analysis of an aliquot in agarose gel, and the rest of the material was kept at −80 °C until needed. Western blot analysis was loaded with 2 different amounts of protein: 20 µg (1) and 40 µg (2).

### Western blot

For western blot analysis, cells were suspended in 50 µl of lysis buffer (50 mM Tris-HCl pH 8.0, 1 mM EDTA, 1 mM DTT, 0.1% Triton X-100, 1% (v/v) NP-40, 1 mM PMSF, 1 μg.ml^-1^ E-64, 1mM sodium fluoride, 1mM sodium orthovanadate) and lysed by sonication. The lysate obtained was centrifuged at 10000 xg for 30 min, and the pellet was resuspended in 20 µl of lysis buffer.

The proteins from total extract, or from isolated chromatin, were loaded onto 15% (w/v) SDS-polyacrylamide gel, resolved by electrophoresis as described previously (26), and electro-transferred to an activated PVDF membrane (Roche). The membranes were blocked with 5% (w/v) non-fat milk suspension in PBS-Tween 0.05% for 2 h, and H3Ser10p was detected using a rabbit aH3Ser10p antibody (ab177218, ABCAM), which was incubated overnight at 4°C. The secondary antibody was incubated for 1 h at room temperature. To develop the signal ECL Plus Western blotting detection system (SuperSignal™ West Femto Maximum Sensitivity Substrate, Thermo Fisher).0

### Flow cytometry

Amastigotes and epimastigotes were first stained with live-dead far red (L34974, INVITROGEN) to detect dead cells and then fixed with the FOXP3 kit (eBioscience FOXP3/Transcription, INVITROGEN).

As indicated by the supplier, after fixation, cells were blocked with blocking solution (perm buffer + 20% fetal serum bovine). They were stained with anti-H3Ser10p (ab177218, ABCAM) 1:300 for 1 h at room temperature, followed by incubation with the secondary anti-Rabbit AF488 (A110008, INVITROGEN) for 1 h at room temperature. Before flow cytometry, cells were stained with 50 μg per sample of Propidium iodide (PI) (SIGMA) to evaluate the cell cycle progression. Data was analyzed with FlowJo VX 0.7 ™, and the Mean fluorescence intensity (MFI) for H3Ser10p signal was recorded.

### Dephosphorylation assays with Lambda protein phosphatase

Epimastigotes were fixed with PFA 4%, after which they were permeabilized with 0.2% Triton-×100 in PBS for 10 min at room temperature. Subsequently, they were incubated with increasing amounts of Lambda protein Phosphatase (P0753S, NEB); during 1 h at 30 °C, in agitation. The samples were subjected to Immunofluorescence and Flow cytometry assays.

## Results

### Nuclear H3Ser10 phosphorylation is restricted to mitotic cells and associates with chromatin in *Trypanosoma cruzi*

Immunofluorescence microscopy of asynchronous epimastigotes cultures revealed a highly restricted distribution of histone H3 serine 10 phosphorylation (H3Ser10p). As shown in Fig. 1A, the H3Ser10p signal was detected exclusively in the nuclei of a small subpopulation of cells corresponding to parasites undergoing mitosis, which were readily identified by the presence of two kinetoplasts (Fig. 1A, inset). In contrast, the vast majority of cells in the culture, corresponding to non-mitotic stages, lacked detectable H3Ser10p staining. Higher-magnification images of merged channels demonstrated a clear colocalization between DAPI and H3Ser10p signals within mitotic nuclei, supporting the nuclear localization of this modification.

**Figure 1.**
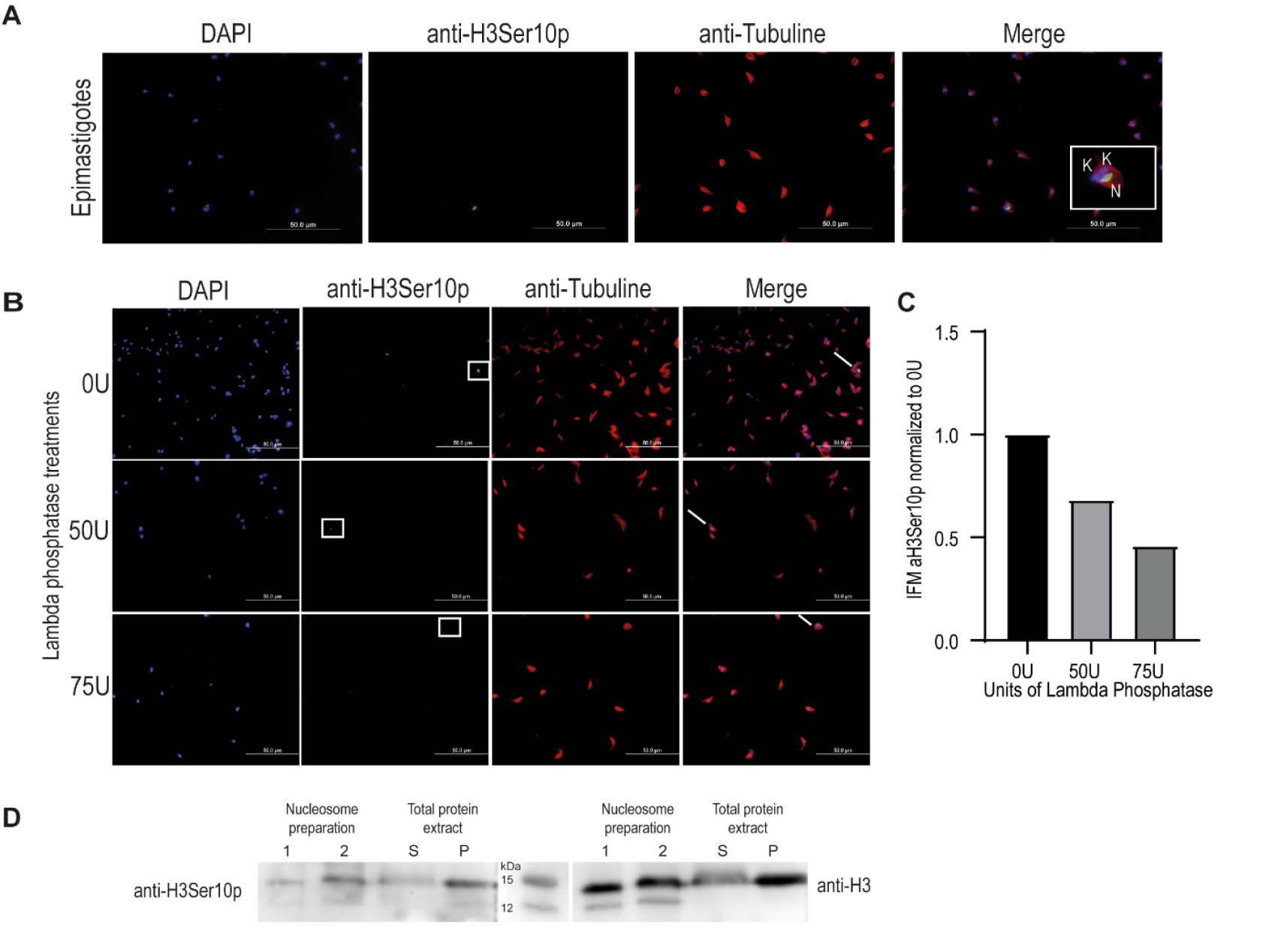
Detection and validation of histone H3 Ser10 phosphorylation in *T. cruzi* epimastigotes. (**A**) Immunofluorescence analysis of asynchronous epimastigote cultures stained with anti-H3Ser10p antibody. DNA from the nucleus (N) and kinetoplast (K) was counterstained with DAPI. Mitotic cells were identified by the presence of two kinetoplasts, as shown in the magnified inset. (**B**) Epimastigotes treated with increasing concentrations of Lambda protein phosphatase for 1 h at 30 °C and stained with anti-H3Ser10p antibody. Loss of signal was observed in a dose-dependent manner. The squares on the panels showing the signal obtained with anti-H3Ser10p and the lines on the Merge panel point to the nuclei in mitosis, where the decrease in signal can be observed as the concentration of Lambda phosphatase increases. (**C**) Quantification of H3Ser10p signal by flow cytometry. Mean fluorescence intensity (MFI) values decrease proportionally with Lambda protein phosphatase concentration. (**D**) Western blot analysis of nucleosome extracts (lane 1: 20 µg and lane 2: 40 µg) and total protein fractions separated into supernatant (S) and pellet (P). The same membrane was probed with anti-H3Ser10p and subsequently re-probed with anti-H3 total to confirm protein size and loading. Anti-tubulin antibody was used to visualize cell morphology, where indicated.

To validate the specificity of the heterologous anti-H3Ser10p antibody in *T. cruzi*, epimastigotes were treated with increasing concentrations of Lambda protein phosphatase before immunofluorescence analysis. As shown in Fig. 1B, Lambda protein phosphatase treatment resulted in a marked and dose-dependent reduction of the nuclear H3Ser10p signal, indicating that antibody recognition depends on the presence of the phosphate group. Importantly, Lambda protein phosphatase treatment did not affect the signal detected with an antibody against total histone H3 (Fig. S1), confirming that the loss of fluorescence was not due to protein degradation or epitope masking.

To quantitatively assess the effect of Lambda protein phosphatase treatment, H3Ser10p levels were measured by flow cytometry. Consistent with the immunofluorescence results, a progressive decrease in the mean fluorescence intensity (MFI) of the H3Ser10p signal was observed as the concentration of Lambda protein phosphatase increased (Fig. 1C), providing independent quantitative support for the phospho-specific nature of the antibody and the modification detected.

Finally, the association of H3Ser10p with chromatin was analyzed by western blot. A single immunoreactive band of approximately 15 kDa was detected using the anti-H3Ser10p antibody in total protein extracts from epimastigotes, as well as in nucleosome preparations (Fig. 1D, left panel). Re-probing the same membrane with an antibody against total histone H3 revealed a single band of comparable molecular weight (Fig. 1D, right panel), supporting the identity of the phosphorylated species. In agreement with a chromatin-associated localization, the H3Ser10p signal was readily observed in the pellet fraction (P). Moreover, analysis of nucleosome core particles obtained by micrococcal nuclease digestion of epimastigotes nuclei confirmed that H3Ser10p is an integral component of chromatin (Fig. 1D, lanes 1 and 2).

### Histone H3 serine 10 phosphorylation is restricted to replicative stages of *Trypanosoma cruzi*

Having established that histone H3 serine 10 phosphorylation is a mitosis-associated, nuclear modification in epimastigotes (Fig. 1), we next investigated whether active cell division is a general requirement for the presence of this modification across different developmental stages of the parasite. To this end, we performed immunofluorescence analysis on three life stages of *T. cruzi* with distinct replication capacities: epimastigotes (replicative stage in the insect vector), intracellular amastigotes harvested at 48 h post-infection (replicative stage in the mammalian host), and trypomastigotes (non-replicative stage).

Consistent with our previous observations, H3Ser10p signal was detected in the nuclei of mitotic epimastigotes (Fig. 2, upper panels). A similar nuclear staining pattern was observed in intracellular amastigotes (Fig. 2, middle panels), where the signal was present in a subset of parasites, in agreement with the asynchronous replication of this stage within host cells.

**Figure 2.**
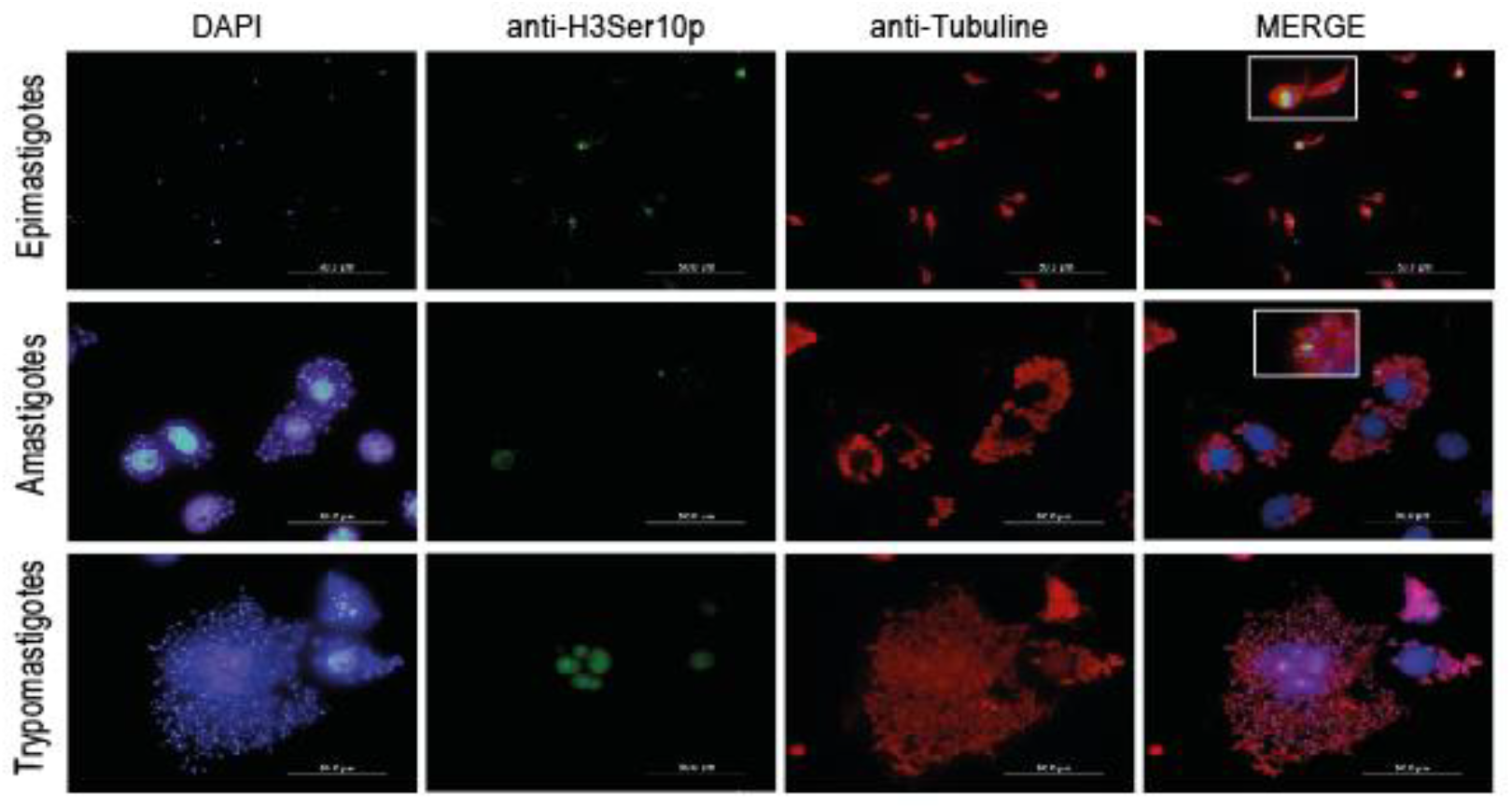
H3Ser10 phosphorylation is detected exclusively in replicative stages of *Trypanosoma cruzi*. Immunofluorescence analysis of epimastigotes (upper panels), intracellular amastigotes (middle panels), and trypomastigotes (lower panels) stained with anti-H3Ser10p antibody (green). Parasite morphology was visualized using anti-tubulin antibody (red), and nuclear and kinetoplast DNA were counterstained with DAPI (blue). Intracellular amastigotes were identified based on morphology and localization within infected host cells. The insets show the overlapping signals of H3Ser10p and DAPI in mitotic cells.

In contrast, no H3Ser10p signal was detected in trypomastigotes (Fig. 2 lower panels), despite clear visualization of nuclear and kinetoplast DNA by DAPI staining and well-preserved parasite morphology as revealed by tubulin labeling. The absence of signal indicates that phosphorylation of histone H3 at serine 10 does not occur in this non-replicative stage.

As a control, total histone H3 was analyzed in parallel and displayed a clear nuclear localization in all life stages examined (Fig. S2), confirming that the lack of H3Ser10p signal in trypomastigotes is not due to reduced histone H3 abundance or impaired antibody accessibility.

Taken together, these results indicate that although histone H3 is present throughout the life cycle of *T. cruzi*, phosphorylation at serine 10 is selectively associated with replicative stages of the parasite and correlates with proliferative capacity.

### H3Ser10 phosphorylation displays dynamic regulation throughout the cell cycle in replicative stages of *Trypanosoma cruzi*

Having shown that histone H3 serine 10 phosphorylation is restricted to replicative stages of the parasite (Figs. 1 and 2), we next investigated the dynamics of this modification throughout the cell cycle. To this end, we analyzed H3Ser10p distribution in individual epimastigotes and intracellular amastigotes by combining immunofluorescence microscopy and flow cytometry-based cell cycle analysis.

The analysis of the dynamics of phosphorylation was facilitated by the utilization of images obtained through immunofluorescence. To determine the cell cycle stage of each parasite, cells were classified based on the number and configuration of nucleus (N), kinetoplast (K), and flagellum (F), a well-established morphological criterion in *T. cruzi*. Briefly, G1 cells display a 1N1K1F configuration, early G2 cells present 1N1K2F, mitotic cells show a 1N2K2F configuration, and post-mitotic cells exhibit a 2N2K2F organization. Using these criteria, we observed that the H3Ser10p signal was detectable at different stages of the cell cycle but was most prominent in cells undergoing mitosis.

Quantification of H3Ser10p fluorescence intensity based on these morphological categories revealed a clear enrichment of the signal in cells displaying the 1N2K2F configuration (Fig. 3A). In asynchronous cultures of both epimastigotes and amastigotes, most cells were found in G1 (1N1K1F or 1N1K, respectively) and lacked detectable H3Ser10p signal. Phosphorylation of H3Ser10 became detectable prior to mitosis (1N1K2F), reached its maximum levels during mitosis (1N2K2F), and declined after cell division, as cells transitioned to the post-mitotic 2N2K2F stage. Notably, all cells classified as 1N2K2F exhibited a discernible H3Ser10p signal.

**Figure 3.**
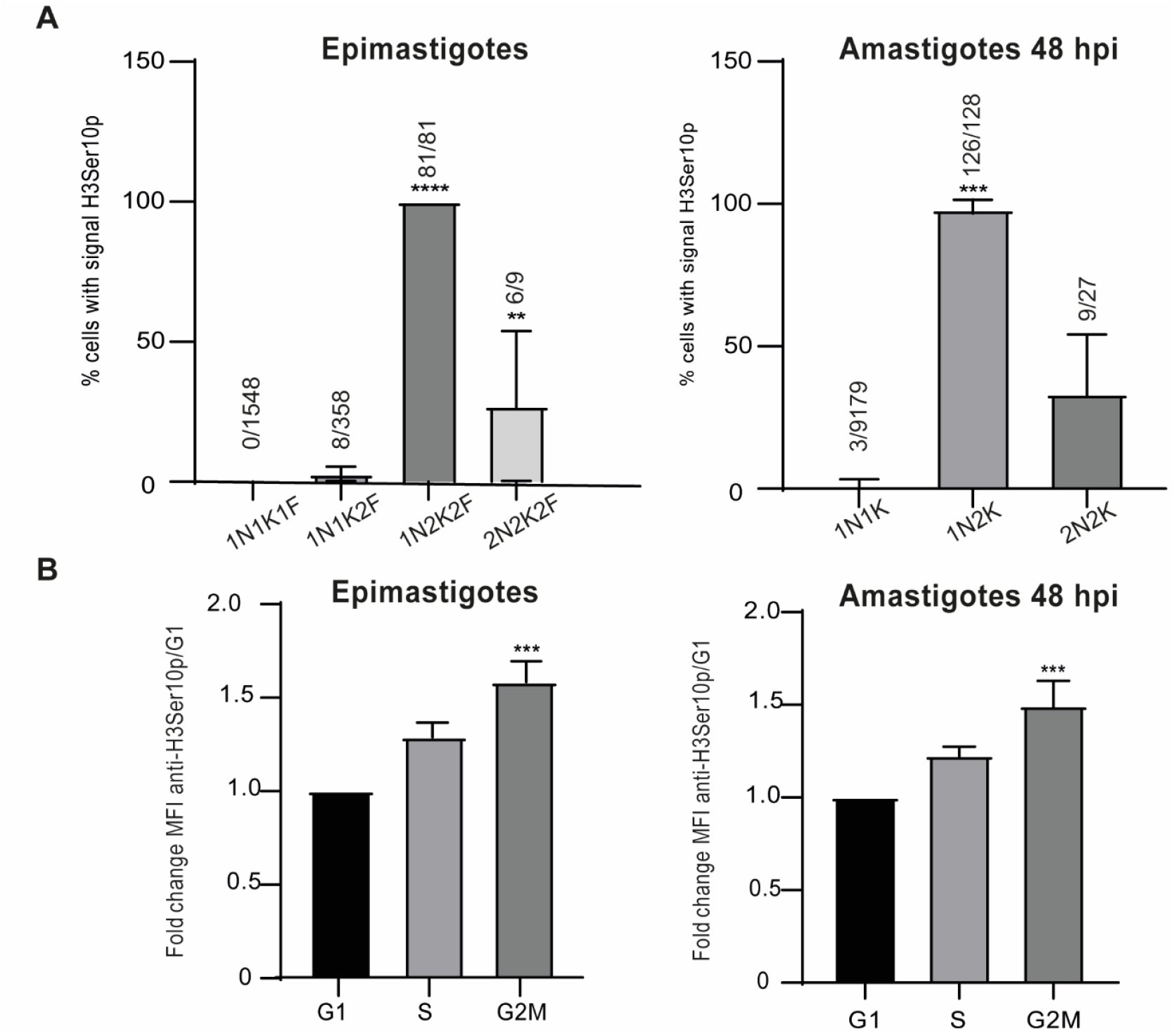
Dynamics of H3Ser10 phosphorylation during the cell cycle in replicative stages of *T. cruzi*. (**A**) Cells were classified according to the number of nucleus (N), kinetoplast (K), and flagellum (F) (1N1K1F, 1N1K2F, 1N2K2F, 2N2K2F), and the ratio of H3Ser10p positive cells to cells at that stage is indicated above each bar. Epimastigotes n = 1996 and amastigotes n = 9334. (**B**) Mean fluorescence intensity (MFI) of anti-H3Ser10p signal in PI-gated cell populations corresponding to different cell cycle stages, normalized to the G1 population, and statistically analyzed using Student’s *t*-test (*** p < 0,001).

To obtain an independent and more quantitative assessment of cell cycle progression, we next analyzed epimastigotes and amastigotes by flow cytometry. Cells were co-stained with propidium iodide (PI) to determine DNA content and with anti-H3Ser10p antibody to assess phosphorylation levels. Flow cytometry analysis allowed us to determine the proportion of cells at each cell cycle stage based on PI fluorescence intensity (Fig. S3).

Finally, we quantified H3Ser10p levels in the different cell cycle stages by measuring the mean fluorescence intensity (MFI) of anti-H3Ser10p within PI-gated populations. When normalized to the G1 population, the H3Ser10p signal was significantly increased in cells in G2/M phases of the cell cycle (Fig. 3B), confirming that phosphorylation of H3Ser10 peaks during mitosis.

To sum up, these results demonstrate that H3Ser10 phosphorylation in *T. cruzi* is a dynamic and tightly regulated process during the cell cycle of replicative stages, reaching maximal levels during mitosis but remaining detectable across multiple phases of cell division.

## Discussion

In eukaryotic cells, cell cycle progression is tightly regulated by multiple factors, among which epigenetic modifications play a central role in ensuring proper transitions between cell cycle phases. One of the best-characterized epigenetic marks associated with mitosis is phosphorylation of histone H3 at serine 10 (1,2). Despite its widespread conservation in eukaryotes, this modification had not been previously detected in trypanosomes. In this work, we provide the first evidence that histone H3 is phosphorylated at serine 10 in *Trypanosoma cruzi*, and we demonstrate that this modification is tightly associated with cell cycle progression and parasite proliferative capacity. Through a combination of immunofluorescence microscopy, biochemical fractionation, and flow cytometry analyses, we show that H3Ser10 phosphorylation is a nuclear, chromatin-associated mark that is restricted to replicative stages and dynamically regulated throughout the cell cycle. Together, these observations establish H3Ser10p as a genuine chromatin modification in this parasite.

The absence of H3Ser10p detection in previous studies of *T. cruzi* is likely attributable to both technical and biological factors. Several earlier works aimed to characterize histone post-translational modifications (PTMs) in this parasite relied primarily on mass spectrometry analyses performed on asynchronous epimastigotes cultures (27–31). Population-based approaches are inherently limited when attempting to detect PTMs that are restricted to specific phases of the cell cycle. In asynchronous cultures, most cells reside in G1, while mitotic cells typically represent only a minor fraction of the population, estimated at approximately 20%. Given that H3Ser10 phosphorylation peaks during mitosis, its signal is likely diluted and masked in bulk analyses. Notably, even a recent study employing synchronized cultures to monitor histone PTMs throughout the cell cycle failed to detect H3Ser10p (31). As we show here, H3Ser10p is only detectable during mitosis or in close temporal proximity to it, time points that were not included in the above-mentioned study. In addition, the transient and potentially unstable nature of this modification suggests that it may be lost during experimental procedures involving extensive washing or harsh sample processing.

In other eukaryotes, H3Ser10 is phosphorylated by Aurora kinase B (AUKB) (6). In *T. brucei*, the homologous enzyme TbAUK1 has been shown to play a key role in cell cycle regulation (20). In *T. cruzi*, TcAUK1 localizes to the extremities of the kinetoplast during interphase, whereas during mitosis it relocates to the nucleus in close association with the mitotic spindle (22). Overexpression of TcAUK1 in epimastigotes results in a delay at the G2/M transition due to impaired kinetoplast duplication (22). Similarly, in *T. brucei*, depletion of TbAUK1 by RNA interference disrupts mitosis, spindle assembly, and chromosome segregation (19,21). More recently, Wiedeman et al. (23) by implementing a novel strategy for a conditional knockout system, concluded that AUK1 and Polo-like kinase (PLK) are essential for proliferation in *T. cruzi* epimastigotes. These observations strongly suggest a conserved role for Aurora kinases in regulating mitotic events in trypanosomatids. Given that histone H3 in *T. cruzi* contains a conserved serine residue at position 10, unlike *T. brucei*, where this residue is absent (19), we hypothesized that H3Ser10 phosphorylation might occur within a very narrow and specific temporal window of the cell cycle.

By employing methods that enable targeted detection of H3Ser10p at the single-cell level, we uncovered the occurrence of this PTM during mitosis. Western blot analyses revealed a single protein band recognized by the anti-H3Ser10p antibody, displaying electrophoretic mobility consistent with histone H3. As expected for a nuclear protein, the signal was evident in the insoluble pellet fraction and was also detected in nucleosome core particle preparations, confirming that H3Ser10p is a bona fide chromatin-associated modification (Fig. 1). Immunofluorescence microscopy further confirmed the nuclear localization of H3Ser10p and revealed its strict association with mitotic cells. Importantly, Lambda protein phosphatase treatment abolished the signal in a dose-dependent manner, demonstrating that phosphorylation of serine 10 is required for antibody recognition.

To further define the temporal dynamics of this modification, we correlated H3Ser10p levels with cell cycle stage using morphological criteria based on the number and position of nucleus, kinetoplast, and flagellum (14,32). While this approach proved robust for epimastigotes, visualization of flagella in amastigotes posed a considerable challenge due to their small size and structural complexity. Therefore, cell cycle staging of amastigotes relied primarily on the number and position of nuclei and kinetoplasts (Fig. 2). Combining immunofluorescence microscopy with flow cytometry allowed us to capture both population-level trends and single-cell resolution. While flow cytometry provides an average estimation of marker abundance across large populations, it lacks the sensitivity to detect subtle differences between individual parasites. In contrast, morphological analysis enabled precise assignment of each cell to a specific cell cycle phase, revealing that H3Ser10 phosphorylation emerges before mitosis, peaks during mitotic progression, and declines after nuclear division (Fig. 3).

Trypanosomes exhibit several unique features when compared to other eukaryotes, including the absence of classical mitotic chromosome condensation. Nevertheless, phosphorylation of H3Ser10 coincides with the maximal level of chromatin compaction reported for trypanosomatids (13). Although this compaction is less pronounced than in higher eukaryotes, it is conceivable that H3Ser10p contributes to the structural reorganization of chromatin required for successful nuclear division. Alternatively, this modification may serve as a platform for the recruitment of additional regulatory factors that promote cell cycle progression.

Beyond its potential structural role, H3Ser10 phosphorylation may also participate in other chromatin-associated processes. In HeLa cells, H3Ser10p has been linked to the formation of R-loops, raising the possibility that a similar association could exist in *T. cruzi*, particularly given the reported accumulation of R-loops during comparable stages of the cell cycle (3). Moreover, in other eukaryotes, H3Ser10p has been shown to functionally interact with H3K9 methylation (33–35). Although H3K9 methylation has not yet been reported in *T. cruzi*, this modification may occur transiently or under specific conditions and therefore remains undetected.

In summary, our study identifies H3Ser10 phosphorylation as a previously unrecognized, cell cycle–regulated chromatin modification in *T. cruzi*. By demonstrating its association with chromatin, its restriction to replicative stages, and its dynamic regulation throughout the cell cycle, we provide new insights into the epigenetic mechanisms that coordinate cell division in this parasite and lay the groundwork for future studies addressing the functional interplay between chromatin modifications and cell cycle control in trypanosomatids. Moreover, nucleosomes and histone PTMs are frequently implicated in developmental changes and affect the pathogenicity of the parasite (36), pointing out their relevance to be addressed as novel therapeutic targets.

## ACKNOWLEGMENTS

We would like to thank Dr. Rafael Argüello and Lic. Rosario Lavignolle for helpful advice on the flow cytometry experiments and for providing us with the anti-H3Ser10p antibody. We would also like to thank Joaquín Iolster for his help with the infection experiments. J.O., S.C.V.L., and G.D.A are members of the Research Career of CONICET. M.d.R.L. is Ph.D. fellow from the same institution, and her PhD thesis is carried out at Departamento de Fisiología, Biología Molecular y Celular, Facultad de Ciencias Exactas y Naturales, Universidad de Buenos Aires.

## Supporting information

**S1 Fig.:**
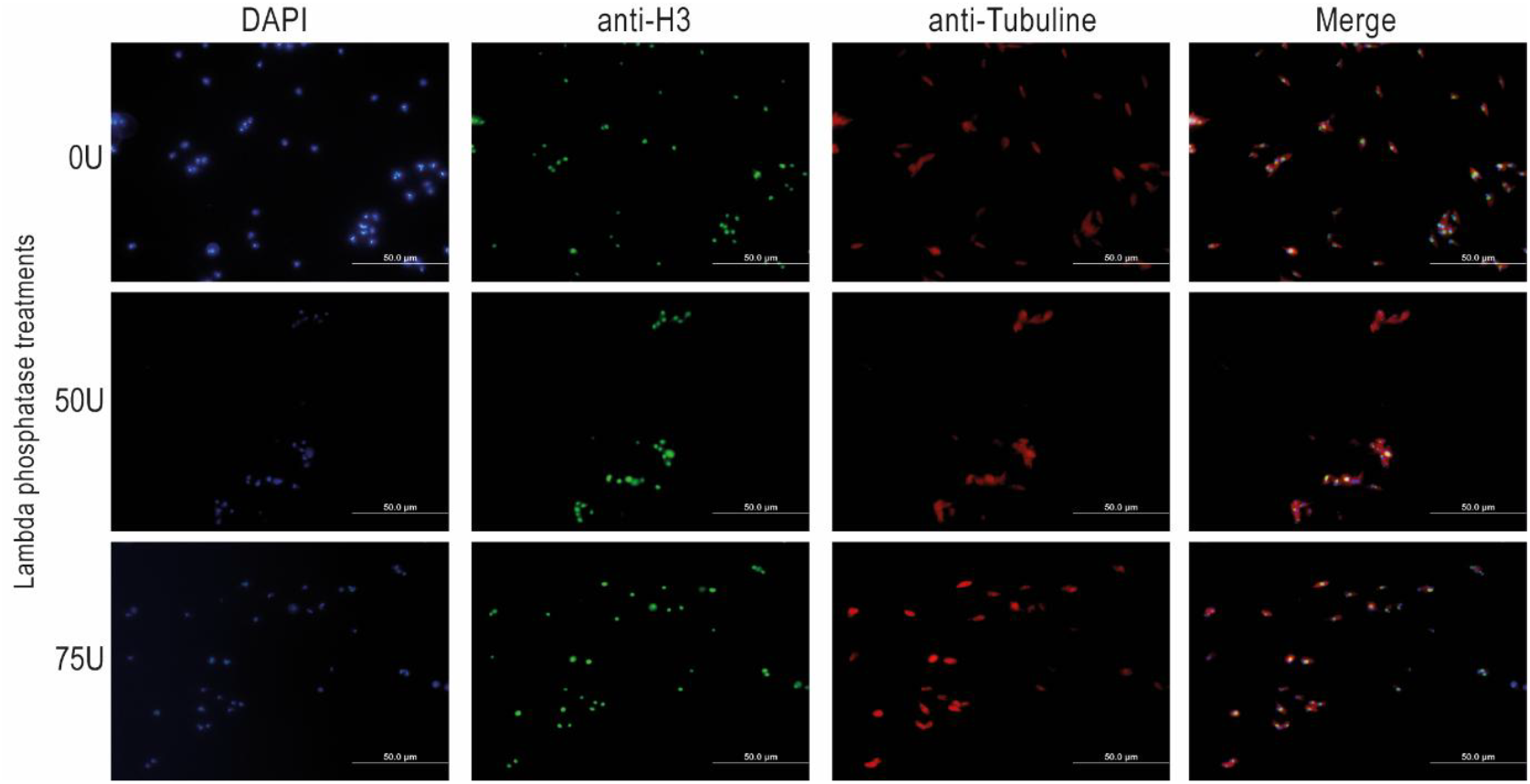
The presence of histone H3 has been detected in every epimastigote treated with Lambda protein phosphatase. Immunofluorescence analysis of epimastigotes stained with anti-H3 (green). Parasite morphology was visualized using anti-tubulin antibody (red), and nuclear and kinetoplast DNA were counterstained with DAPI (blue).

**S2 Fig.:**
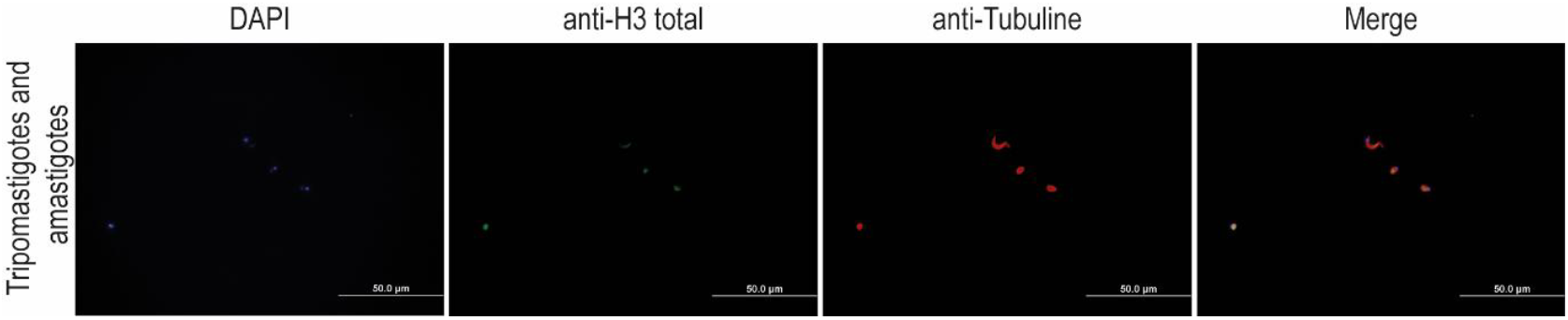
The presence of histone H3 has been observed in both trypomastigotes and amastigotes. Immunofluorescence analysis of trypomastigotes and amastigotes stained with anti-H3 (green). Parasite morphology was visualized using anti-tubulin antibody (red), and nuclear and kinetoplast DNA were counterstained with DAPI (blue). Histone H3 is present throughout all stages of the life cycle of *T. cruzi*.

**S3 Fig.:**
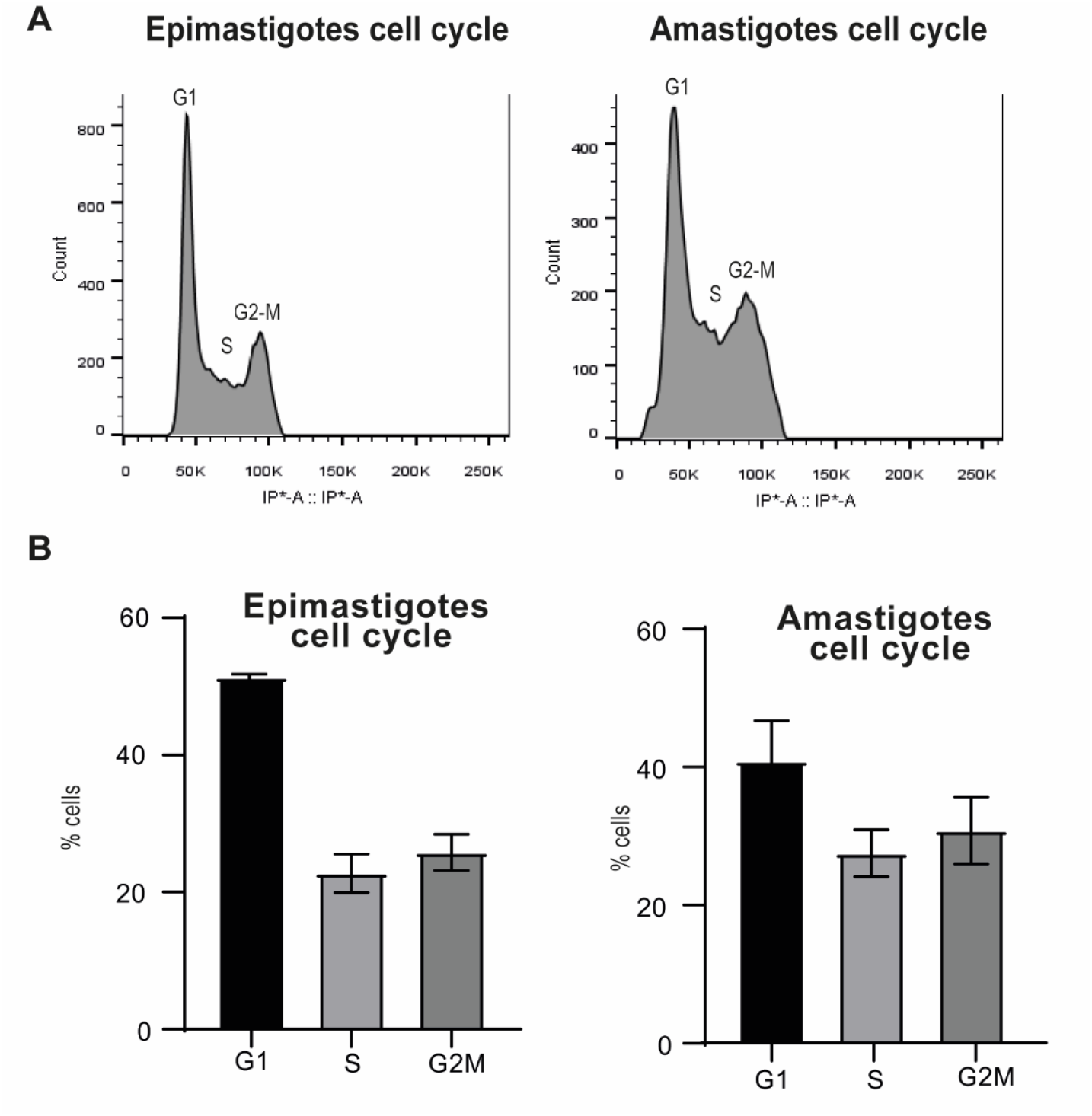
Cell cycle measurements for flow cytometry. The subpopulations of epimastigotes and amastigotes were distinguished by means of flow cytometry, with this distinction being made according to the quantity of DNA and, consequently, the cell cycle stage. **(A)** Histogram. **(B)** % of cells in each stage using the MFI of PI

## References

1. Hendzel MJ, Wei Y, Mancini MA, Van Hooser A, Ranalli T, Brinkley BR, et al. Mitosis-specific phosphorylation of histone H3 initiates primarily within pericentromeric heterochromatin during G2 and spreads in an ordered fashion coincident with mitotic chromosome condensation. Chromosoma. 1997 Nov 19;106(6):348–60.

2. Hans F, Dimitrov S. Histone H3 phosphorylation and cell division. Oncogene. 2001 May 28;20(24):3021–7.

3. Castellano-Pozo M, Santos-Pereira JM, Rondón AG, Barroso S, Andújar E, Pérez-Alegre M, et al. R Loops Are Linked to Histone H3 S10 Phosphorylation and Chromatin Condensation. Mol Cell. 2013 Nov;52(4):583–90.

4. Ruchaud S, Carmena M, Earnshaw WC. Chromosomal passengers: conducting cell division. Nat Rev Mol Cell Biol. 2007 Oct;8(10):798–812.

5. Van De Werken C, Avo Santos M, Laven JSE, Eleveld C, Fauser BCJM, Lens SMA, et al. Chromosome segregation regulation in human zygotes: altered mitotic histone phosphorylation dynamics underlying centromeric targeting of the chromosomal passenger complex. Hum Reprod. 2015 Oct;30(10):2275–91.

6. Crosio C, Fimia GM, Loury R, Kimura M, Okano Y, Zhou H, et al. Mitotic Phosphorylation of Histone H3: Spatio-Temporal Regulation by Mammalian Aurora Kinases. Mol Cell Biol. 2002 Feb 1;22(3):874–85.

7. Giet R, Prigent C. The Xenopus laevis Aurora/Ip11p-Related Kinase pEg2 Participates in the Stability of the Bipolar Mitotic Spindle. Exp Cell Res. 2000 Jul;258(1):145–51.

8. Chadee DN, Hendzel MJ, Tylipski CP, Allis CD, Bazett-Jones DP, Wright JA, et al. Increased Ser-10 Phosphorylation of Histone H3 in Mitogen-stimulated and Oncogene-transformed Mouse Fibroblasts. J Biol Chem. 1999 Aug;274(35):24914– 20.

9. Hsu JY, Sun ZW, Li X, Reuben M, Tatchell K, Bishop DK, et al. Mitotic Phosphorylation of Histone H3 Is Governed by Ipl1/aurora Kinase and Glc7/PP1 Phosphatase in Budding Yeast and Nematodes. Cell. 2000 Aug;102(3):279–91.

10. Juan G, Traganos F, James WM, Ray JM, Roberge M, Sauve DM, et al. Histone H3 phosphorylation and expression of cyclins A and B1 measured in individual cells during their progression through G2 and mitosis. Cytometry. 1998 Jun 1;32(2):71–7.

11. Jacobberger JW, Frisa PS, Sramkoski RM, Stefan T, Shults KE, Soni DV. A new biomarker for mitotic cells. Cytometry A. 2008 Jan;73A(1):5–15.

12. Ogbadoyi E, Ersfeld K, Robinson D, Sherwin T, Gull K. Architecture of the Trypanosoma brucei nucleus during interphase and mitosis. Chromosoma. 2000 Mar 28;108(8):501–13.

13. Spadiliero B, Nicolini C, Mascetti G, Henríquez D, Vergani L. Chromatin of Trypanosoma cruzi : In situ analysis revealed its unusual structure and nuclear organization. J Cell Biochem. 2002 Jan;85(4):798–808.

14. Elias M, Dacunha J, Defaria F, Mortara R, Freymuller E, Schenkman S. Morphological Events during the Trypanosoma cruzi Cell Cycle. Protist. 2007 Apr 18;158(2):147–57.

15. Shlomai J. The Structure and Replication of Kinetoplast DNA.

16. Gluenz E, Povelones ML, Englund PT, Gull K. The Kinetoplast Duplication Cycle in Trypanosoma brucei Is Orchestrated by Cytoskeleton-Mediated Cell Morphogenesis. Mol Cell Biol. 2011 Mar 1;31(5):1012–21.

17. Hammarton TC. Cell cycle regulation in Trypanosoma brucei. Mol Biochem Parasitol. 2007 May;153(1):1–8.

18. Nunes VS, Moretti NS, Da Silva MS, Elias MC, Janzen CJ, Schenkman S. Trimethylation of histone H3K76 by Dot1B enhances cell cycle progression after mitosis in Trypanosoma cruzi. Biochim Biophys Acta BBA - Mol Cell Res. 2020 Jul;1867(7):118694.

19. Li Z, Wang CC. Changing Roles of Aurora-B Kinase in Two Life Cycle Stages of Trypanosoma brucei. Eukaryot Cell. 2006 Jul;5(7):1026–35.

20. Li Z, Umeyama T, Wang CC. The Aurora Kinase in Trypanosoma brucei Plays Distinctive Roles in Metaphase-Anaphase Transition and Cytokinetic Initiation. Ullu E, editor. PLoS Pathog. 2009 Sep 11;5(9):e1000575.

21. Tu X, Kumar P, Li Z, Wang CC. An Aurora Kinase Homologue Is Involved in Regulating Both Mitosis and Cytokinesis in Trypanosoma brucei. J Biol Chem. 2006 Apr;281(14):9677–87.

22. Fassolari M, Alonso GD. Aurora kinase protein family in Trypanosoma cruzi: Novel role of an AUK-B homologue in kinetoplast replication. Buscaglia CA, editor. PLoS Negl Trop Dis. 2019 Mar 21;13(3):e0007256.

23. Wiedeman J, Harrison R, Etheridge RD. A limitation lifted: A conditional knockdown system reveals essential roles for Polo-like kinase and Aurora kinase 1 in Trypanosoma cruzi cell division. Proc Natl Acad Sci. 2025 Feb 25;122(8):e2416009122.

24. Ocampo J, Carena S, López MDR, Vela VS, Zambrano Siri RT, Balestra SA, et al. Trypanosomatid histones: the building blocks of the epigenetic code of highly divergent eukaryotes. Biochem J. 2025 Mar 18;482(06):325–40.

25. Marques A, Nakayasu E, Almeida I. extracellular and intracellular amastigotes of Trypanosoma cruzi from mammalian host-infected cells. Protoc Exch.

26. Laemmli UK. Cleavage of structural proteins during the assembly of the head of bacteriophage T4. Nature. 1970 Aug 15;227(5259):680–5.

27. De Jesus TCL, Nunes VS, Lopes MDC, Martil DE, Iwai LK, Moretti NS, et al. Chromatin Proteomics Reveals Variable Histone Modifications during the Life Cycle of Trypanosoma cruzi. J Proteome Res. 2016 Jun 3;15(6):2039–51.

28. Picchi GFA, Zulkievicz V, Krieger MA, Zanchin NT, Goldenberg S, De Godoy LMF. Post-translational Modifications of Trypanosoma cruzi Canonical and Variant Histones. J Proteome Res. 2017 Mar 3;16(3):1167–79.

29. De Lima LP, Poubel SB, Yuan ZF, Rosón JN, Vitorino FNDL, Holetz FB, et al. Improvements on the quantitative analysis of Trypanosoma cruzi histone post translational modifications: Study of changes in epigenetic marks through the parasite’s metacyclogenesis and life cycle. J Proteomics. 2020 Aug;225:103847.

30. De Almeida RF, Fernandes M, De Godoy LMF. An updated map of Trypanosoma cruzi histone post-translational modifications. Sci Data. 2021 Mar 25;8(1):93.

31. Menezes APJ, Silber AM, Elias MC, Da Cunha JPC. Trypanosoma cruzi cell cycle progression exhibits minimal variation in histone PTMs with unique histone H4 acetylation pattern. J Proteomics. 2025 May;315:105413.

32. Elias MC, Nardelli SC, Schenkman S. Chromatin and Nuclear Organization in Trypanosoma Cruzi. Future Microbiol. 2009 Oct;4(8):1065–74.

33. Fischle W, Tseng BS, Dormann HL, Ueberheide BM, Garcia BA, Shabanowitz J, et al. Regulation of HP1–chromatin binding by histone H3 methylation and phosphorylation. Nature. 2005 Dec;438(7071):1116–22.

34. Hirota T, Lipp JJ, Toh BH, Peters JM. Histone H3 serine 10 phosphorylation by Aurora B causes HP1 dissociation from heterochromatin. Nature. 2005 Dec;438(7071):1176–80.

35. Mallm JP, Rippe K. Aurora Kinase B Regulates Telomerase Activity via a Centromeric RNA in Stem Cells. Cell Rep. 2015 Jun;11(10):1667–78.

36. Deák G, Wilson MD. Parasite nucleosomes: Chromatin dynamics rewired. Lukeš J, editor. PLOS Pathog. 2025 Dec 15;21(12):e1013781.

